# Pan-genome evolution and its association with divergence of metabolic functions in Bifidobacterium genus

**DOI:** 10.1101/2022.04.05.487182

**Authors:** Sushanta Deb

**Affiliations:** Tripura University, Suryamaninagar, Tripura (W)-799022, India; All India Institute of Medical Sciences (AIIMS), New Delhi 110029, India

**Keywords:** Genome evolution, Horizontal Gene Transfer, Bifidobacteria, Pangenome and Core genome

## Abstract

Many works have been performed to characterize the genomic evolution and diversity among type species of the Bifidobacterium genus due to its health-promoting effect on their host. However, those studies were mainly based on species-level taxonomic resolution, adaptation, and characterization of carbohydrate metabolic features of the bifidobacterial species. Here, a comprehensive analysis of the type strain genome unveils the association of pan-genome evolution with the divergence of metabolic function of the Bifidobacterium genus. This study also has demonstrated that the horizontal gene transfer and genome expansion and reduction events in the evolutionary history influencing the diversity of metabolic functions of bifidobacterium genus. Furthermore, the genome-based search of probiotic traits among all the available bifidobacterial type strains gives hints on type species, that could confer health benefits to nutrient-deficient individuals. Altogether, the present study provides insight into the developments of genomic evolution, functional divergence, and potential probiotic type species of the Bifidobacterium genus.

## Introduction

The members of the Bifidobacterium genera are among the most dominant probiotic bacteria in human gut microbiota, adopts multiple modes to confer health benefits to the host, such as hindering inflammatory reactions [1], synthesis of nutrient [2], strengthening the immune system [3] and competing with pathogenic microbes [4]. A number of bifidobacterial species were reported to have health-promoting effects on a wide range of hosts [5]. Genome-based comparative analysis has always been exploited to decipher the changes in evolutionary traits of any bacterial taxonomic group and provides insight into the genetic factors responsible for bacterial adaptation to a specific ecological niche [6]. Several studies on Phylogenomic and comparative genomic approaches have been reported targeting the entire genus [7–10] and individual probiotic members of Bifidobacterium [11–13], those studies were confined to the role of polysaccharides for adaptation [7, 8], and taxonomic demarcation of bifidobacterial species [9, 10]. The diverse habitats of closely related bacterial species can result in variability in their genetic contents and physiological functions [14, 15]. The variable genetic content (pan-genome) emerges across the isolates of the same taxonomic group mainly due to the frequent gene gain-loss events in bacterial genomes [16, 17]. Several earlier studies on bifidobacteria explained genome evolution and dynamics of gene content. [18–20]; however, little information about the functional divergence and probiotic potential of all bifidobacterial type strains is available to date. In recent time, this genera comprised 86 validly published novel species (http://www.bacterio.net/bifidobacterium.html; accessed 18^th^ Aug 2021). However, only 75 novel species genomes are publicly available in the NCBI database.

This study aimed to correlate the pan-genome evolution with the functional divergence of the type strains of the Bifidobacterium genus. The present study also describes the gene content dynamics and horizontal gene transfer events to correlate the genome evolution and divergence of metabolic function of the Bifidobacterium genus. Additionally, Core-genome based phylogeny has found the genomic footprints of host-specific selection pressure operating on the bifidobacterial species. Furthermore, genome-based screening for the health-promoting potential of all the analyzed type species gives insight into unexplored probiotic candidates of the Bifidobacterium genus.

## Methods

### Data retrieval and quality assesment

The genome sequences of 75 type strains of bifidobacterial species are defined by LPSN database (accessed 18th Aug 2021). A total of 75 complete sequenced type strain genomes of the bifidobacterial genus were obtained from NCBI databases (Supplimentary Table-1). Contamination and completeness of genomes retrieved from the NCBI database were estimated using CheckM v1.1.3 [21]. Incomplete genome sequence (< 90%) of bifidobacterium commune discarded from the downstream analysis. The average nucleotide identity (ANIb) calculations were performed using pyani.py (https://github.com/widdowquinn/pyani).

### Core genome and pan-genome phylogeny construction

A BlastP search was performed with an e-value cutoff of 1.0 e-15 among the genomes of 74 type strains downloaded from the NCBI database to construct single-copy core gene families. The hits were selected with ≥50% and ≥70% of alignment coverage, as previously described in Staphylococcus and Acinetobacter genus [22, 23]. The recombinant genes were removed from the core gene families with the help of PhiTest feature implemented in PhiPack program [24].

The non-recombinant core genes were concatenated to reconstruct a maximum-likelihood tree with the generalized time-reversible (GTR) model using the RAxML tool [25]. The pan-genome was determined with the help of the Ortho Markov Cluster (OMCL) [26] and Clusters of Orthologous Groups triangles (COGtriangles) algorithm implemented in the GET_HOMOLOGOUS program [27], as previously described in a case study on pIncA/C Plasmids [28]. Two auxiliary Perl-script, compare_clusters.pl and parse_pangenome_matrix.pl, were used to generate the pan-genome matrix. This matrix file was subsequently used as an input for estimate_pangenome_phylogenies.sh program to generate the maximum-likelihood (ML) pan-genome phylogenetic tree [29]. The branch order similarity between core and pan-genome tree was evaluated using TreeCmp [30], calculating two scores, i.e., normalized matching-cluster (nMC) score, and normalized Robinson-Foulds (nRF) score. The phylogenetic trees were visualized using the Interactive Tree Of Life (iTOL) v5 program [31].

### COG functional annotation

The clusters of orthologous groups (COG) functional annotation of coding sequences was performed using the Perl script CDD2COG.pl v0.2 (https://github.com/aleimba/bac-genomics-scripts/tree/master/cdd2cog). Potential probiotic genes among the bifidobacterial genomes were identified based on COG functional annotation. The heatmap representing the distribution of genes in different COG functional categories was generated using the manhattan distance and average clustering method embedded in the heatmap2 function of the gplots package [32] in R program[33].

### Genome dynamics and evolution

To obtain a larger view into the evolutionary dynamics of gene families, ancestral reconstruction based evolutionary history of 74 bifidobacterial species were deduced using the Count program based on the Dollo parsimony method with automated rate optimization criterion [34, 35]. Horizontal gene transfer events were analyzed using HGTector [36] software, which follows blastP based sequence similarity search (E value of 1e-10, percentage identity of 40%, and query coverage of 70%) with the help of DIAMOND v2.0.4 [37] toll against nonredundant RefSeq prokaryotic protein database (October 2019). During the HGTector run, Bifidobacterium was set as the self group, and Bifidobacteriaceae was set as the close group. The potential horizontal genes among 74 bifidobacterial genomes were extracted from the output files of HGTector pipeline.

## Result

### General genomic features

Seventy-five type strain genomes from the bifidobacterium genus were obtained from the NCBI database. The genomes were sorted according to their source, human, monkey, lemur, rabbit, rodent, sewage, and fermented milk isolate. Overall, the genome sizes and GC contents ranged from 1.51 to 3.25 Mb and 66.64 to 50.37 percent, respectively (supplimentary Table-1). The ANI values were calculated to estimate the genomic similarity among type strains of the bifidobacterium genus. A phylogenetic tree was constructed based on 188 core-gene families shared among 74 (B.commune excluded for poor quality genome) bifidobacterium type strains to demonstrate the evolutionary relationships among the isolates of different niches. Type strains isolated from various hosts were clustered in distinct clades of the phylogenetic tree. The ANI based genomic distance suggested all the type strain genomes analyzed in this study follows the defined value (95-96%) for species demarcation [38] and are taxonomically distinct at the species level.

### Core and Pan-genome Phylogeny

We analyzed the core and pan-genome among 74 Bifidobacterium type strains to evaluate their genomic diversity. To draw the inference on the evolutionary relationship among type strains, a core genome tree was constructed using concatenated nucleotide sequences of 188 single copy ortholog genes present in all 74 type strains. Core genome phylogeny exhibits the number of distinct genome clusters, indicating genetic similarity varies with different groups of type species of the bifidobacterium genus (figure-1). Additionally, a pan-genome tree was constructed and compared with the core tree to evaluate the correlation between phylogeny and the genetic variability of this taxonomic group. The pan-genome analysis showed the existence of 18084 pan-gene families among the type species of bifidobacterium genus. Comparison between two trees unveiled inconsistency in branching order and phylogenetic position of type strain genomes (figure-2). To assess the range of discrepancy in ancestral placement and topology of core and pan-genome tree, two values such as normalized matching-cluster (nMC) [30] and normalized Robinson-Foulds (nRF) [39] were calculated. The values ranging from 0 to 1, if the value deviate from 0 to 1, indicate a lack of conformity between the two trees. In the present study, observed values for nMC and nRF were 0.31 and 0.43, respectively, which represents that the lineage history of the pan-genome is not uniform with the evolution of core genes among the members of the Bifidobacterium genus.

**figure 1:**
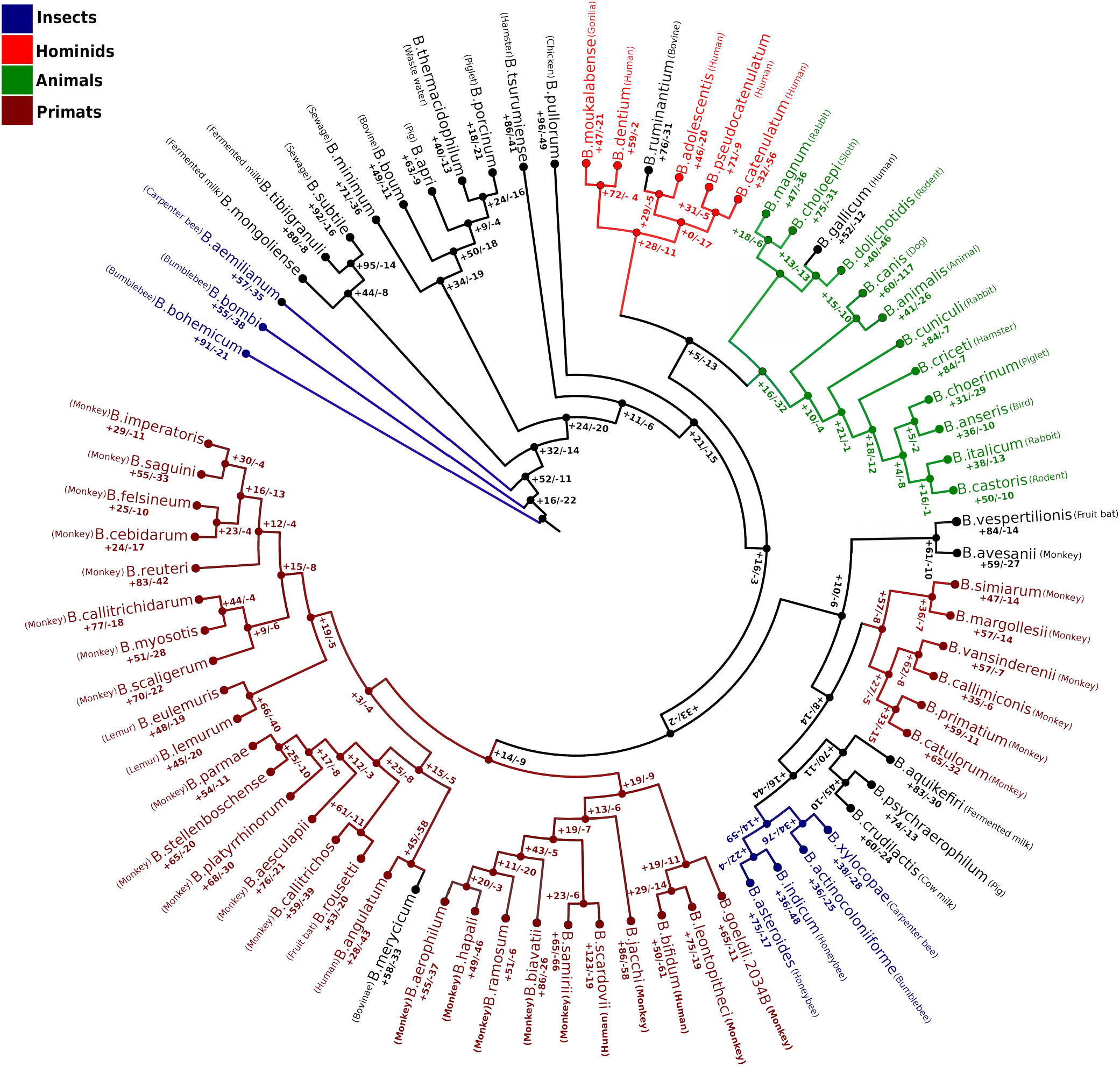
Evolutionary history of type species annotated with isolation source, estimated gene gain-loss event across the lineage of bifidobacterial evolution.

**figure 2:**
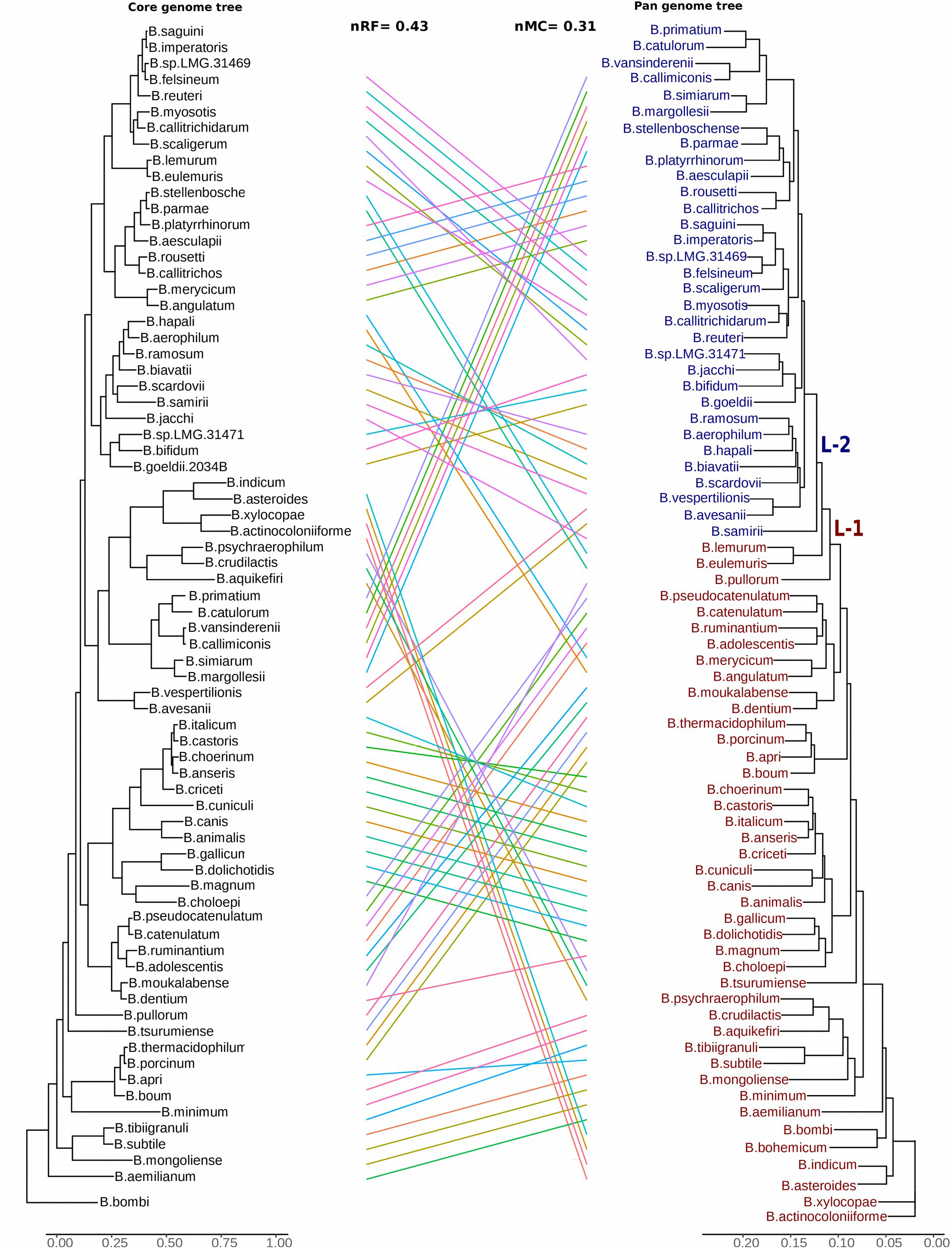
Combined Core and Pan-genomes based Phylogenetic tree showing a diverse evolutionary pattern of the variable genetic content of bifidobacterial type species.

### COG functional profile

The COG based functional profile was obtained to understand the functional variability in the genetic repertoire of genus bifidobacterium. To discriminate the distinct functional profile of type strains, a heat map was generated to visualize the distribution of COG functional categories across the genus. Surprisingly, hierarchical clustering based on COG functions divides the type strain genomes into two clusters named L1 and L2 (figure-3), which is according to the clustering pattern of the pan-genome phylogenetic tree (figure-2). The heat map demonstrated a higher percentage of genes of different COG functions present in the L2 cluster than that of the L1 cluster (figure-3). In addition, the functional profile of core genes has also been investigated to represent the variability in relative COG functional abundance of core genes (Supplimentary figure-1). However, hierarchical clustering based on core gene function has not shown any distinct clustering pattern of type strain genomes (suppliment figure-1).

**figure 3:**
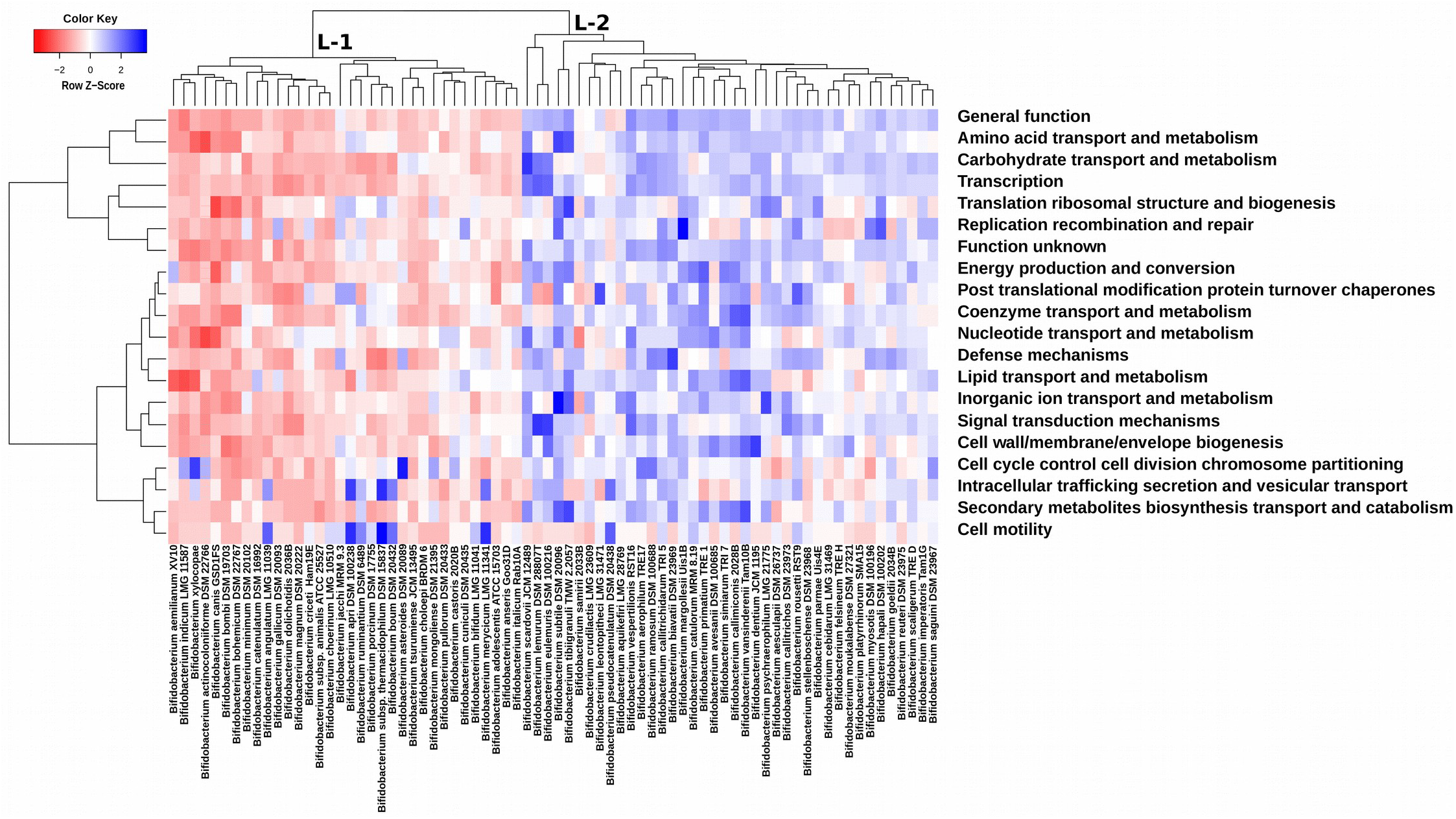
COG frequency heatmap representing two clusters (L-1 and L-2) of type species based on the profile of metabolic function.

### Evolutionary history, horizontal genes and host specific phylogenetic groups in bifidobacterium genus

To investigate the gene gain and loss events throughout the evolution of each ancestor and descendants, we performed the ancestral reconstruction and gene content analysis for the 74 type strain of the Bifidobacterium genus. Gene gain-loss phenomenon improves adaptive capability through the course of evolution and diversifies prokaryotes’ metabolic functions [40, 41]. The phylogenomic tree showed that the frequency of gain-loss events is higher in descendants than in common ancestors (fig). In addition, it has also been observed that the occurrence of gene gain was greater than that of gene loss in the evolutionary history of the Bifidobacterium genus. Furthermore, the gene loss count in B. Canis (117 lost genes) was significantly higher than any other bifidobacterial species, suggesting this species might have evolved under greater environmental selection pressure.

Horizontal genes are the major driving force influencing the genetic diversity and bacterial speciation in habitats [42]. The analysis of horizontally transferred genes for each type strain revealed that a major portion of the genetic repertoire in Bifidobacterium genus was acquired horizontally from the other isolates of the habitat (supplimentary Table-2). The number of horizontally transferred genes in type strain genomes are listed in Table (supplimentary Table-2).

The core genome phylogenetic tree has shown a possible association between evolutionary relations and isolation source of type species. The isolated genomes from various host types such as insects, animals, primates and hominids were clustered into a distinct phyletic group (clade) based on their isolation source (figure-1).

### Distribution of probiotic traits across the bifidobacterium genus

An isolate must confront the unfavorable gastrointestinal (GI) environment, such as acid, bile salt concentrations, heat, and osmotic pressure to confer the probiotic function to the host [43]. Functional analysis has identified the distribution of genes related to stress tolerance response, such as cobalt transporter (COG0619), heat shock molecular chaperone (COG0576), universal stress protein (COG0589), and methionine sulfoxide reductase (COG0225) [44]. Earlier reports suggested that bacteria contribute to vitamin synthesis in the human gut and provide health benefits to the host [45, 46]. Furthermore, this study also demonstrated the vitamin synthesis capability of the type strains belonging to the Bifidobacterium genus. It has been identified several potential genes involved in Thiamine, Riboflavin, Biotin and Folate metabolism were distributed among the Bifidobacterium species. Adhesion capability is crucial for probiotic isolates; it competes with enteropathogens for attachment in the Gastrointestinal tract [47]. Studies have reported that enolase (COG0148) and Triosephosphate isomerase (COG0149) has a potential role in cell surface adherence [48, 49]. Functional annotation of bifidobacterial genomes has shown the distribution of adherence associated genes among the type species of the Bifidobacterium genus. The genes related to probiotic functions considered in this study were listed in the table S3 (Supplimentary Table-3).

## Discussion

The nRF and nMC scores for pan and core-genome evolution tree were 0.43 and 0.31, respectively, which implies that phyletic relation of pan gene families was not completely congruent with the core genes of bifidobacterium genus. This inconsistency between two phylogenetic group can be explained by expansion of accesory gene pool and acquisition of exogenous genetic material through the horizontal transfer events within the genus. The occurence of large number of pan gene families (18084 gene) among the type species of bifidobacterium may be due to the wide range of hosts and habitats of this genus. This study demonstrated that presence of variable pan-genes might have been playing a crucial role in evolution of type species of bifidobacterial genus and made them competent to survive in various environmental pressures. Out of 34 members in L2 clade of pangenome phylogenetic tree 32 were consistent with the type species of L2 clade in COG based functional heatmap. This congruence in taxa distribution between pan genome tree and COG frequency heatmap, support the notion that evolution of pangenomic content (variable portion of genomes) may have led to the functional enrichment within the species of that phylogenetic group of bifidobacterium genus. The major variation in COG profile was mainly contributed to the emergence of two functionally similar groups on heirarcheal clusturing of type species. Furthermore, COG functional analysis on core genes have revealed an higher abundance of genes in three categories i.e 1. Translation ribosomal structure and biogenesis followed by the 2. General function and 3. Posttranslational modification protein turnover chaperones (Supplimentary figure2).

A relationship between the type species phylogeny and the hosts of bifidobacteria were observed, isolates from the hosts of same taxonomic group were clustered into a single clade. This observation suggesting that a host specific selection pressure might have been operating on the core-genes of bifidobacterium genus.Nevertheless, several strains (colured in black) showed unbalanced distribution with no clear patterns in host based phylogenetic clading as they are distributed in different clades of the core genes evolutionary tree. This indicates that cross species transmission or host jumping and other routes of spreading have widen the range of ecologilcal niches for bifidobacterial species.

Bottacini et al. [50] have proposed that insects are the most ancient host of bifidobacteria. However, in a subsequent analysis it has been speculated B. psychraerophilum and B. Minimum emerged from the most ancient common ancestor in bifidobacterial evolution [19]. Although, in this study it is observed that strains isolated from bees descended from the last common ancestor (LCA) of bifidobacterium parellel with the finding of Bottacini et al. [50]. Moreover, present study also demonstared that isolates from human and other primates dominates the younger lineage of evolutionary tree, supporting the hypothesis that spreading of bifidobacterial strains occured from nonprimate animals to primates like humans and monkeys [19].

Almost, every single nodes (internal and external) of evolutionary tree exhibits gene expansion and contraction events, which indicates a stringent genome streamlining strategy and dynamic evolution of bifidobacterium species (figure-1). Bifidobacterium experienced numerous gene gain events that might have yield myriad of functional diversity along the course of evolutionary process [51]. As expected, consistent with the gene gain events, a massive frequency of horizontal gene transfer (HGT) events were also observed in the members of bfidobacterium group (supplimentary Table-2).

The putative taxa from which genes were horizontaly transfered in bifidobacterium were identified as Actinobacteria, Bacillus, Gammaproteobacteria, Clostridia and Alphaproteobacteria [7]. It is reported, all the putative taxa identified as donor for HGT events in bifidobacterium are predominant in the gut environment [52]. Present analysis have revealed that amongst all the type strains Bifidobacterium scardovii aquired the highest number of genes (669) from the other microbes (supplimentary Table-2). Several earlier studies have reported HGT as a major influencing factor transforming the non-pathogenic strains into potent pathogenic isolate [53–55]. Most of the bifidobacterial species are from the normal gut flora of human and animal, they are mainly non-pathogenic in nature and confers probiotic benefits to host [56]. Exceptionally, Bifidobacterium scardovii was found to be associated with urinary tract infection (UTI) and considered as an potential pathogen for UTIs infection [57]. Considering that it can be speculated, evolution of pathogenic traits may likely be due to the higher frequency of HGT events in B. Scardovii. In addition, a close look into the number of HGT genes in each type strains of genus bifidobacteria, categorized the type species into two groups low HGT frequency and high HGT frequency group highlighted in yellow and red respectively in supplimentary Table-2. Interestingly, the species with low HGT frequency and high HGT frequency are grouped together and are in congruence with the bifidobacterial species group of L1 and L2 cluster in COG based functional heatmap respectively (figure2). This observation indicates that high frequency HGT events have made the species from L2 clusture functionally more enriched. Looking into the another aspect, consistency in species distribution between pan genome phylogenetic group and L2 clusture of functional heatmap, suggesting a possible link of HGT event with the pangenome evolution and functional diversity in genus bifidobacterium. Several members of bifidobacterium genus are involved in providing health benifits to the host [58–60]. This makes the bifidobacterium a potential genus that may contain newly discovered species with unexplored probiotic traits. This study attempted to gain genome based preleminary insight on the probable probiotic features of all the available type strains of this genus. As shown in figure-4, there is hardly any variation in relative abundance of genes associated with stress tolerence and adhesions, however a substantial variation has been observed in abundancy of the genes related to vitamin synthesis (Thiamine, riboflavin, folate and biotin) across the different type species of bfidobacterium genus (figure-4). Few type species have shown a higher abundance of gene involved in riboflavine and biotin metabolism. Riboflavin metabolism is dominant in asteroids, bombi and xylocopae while genes for biotin metabolism are mainly profund in B. adolescentis, B. catenulatum, B. dentium,B. Merycicum and B. tibiigranuli (figure-4). Higher abundancy of genes involved for vitamin metabolism in the above mention type species implying their capabilty to be a succesful probiotic candidate to the nutrient deficiate individuals.

**figure 4:**
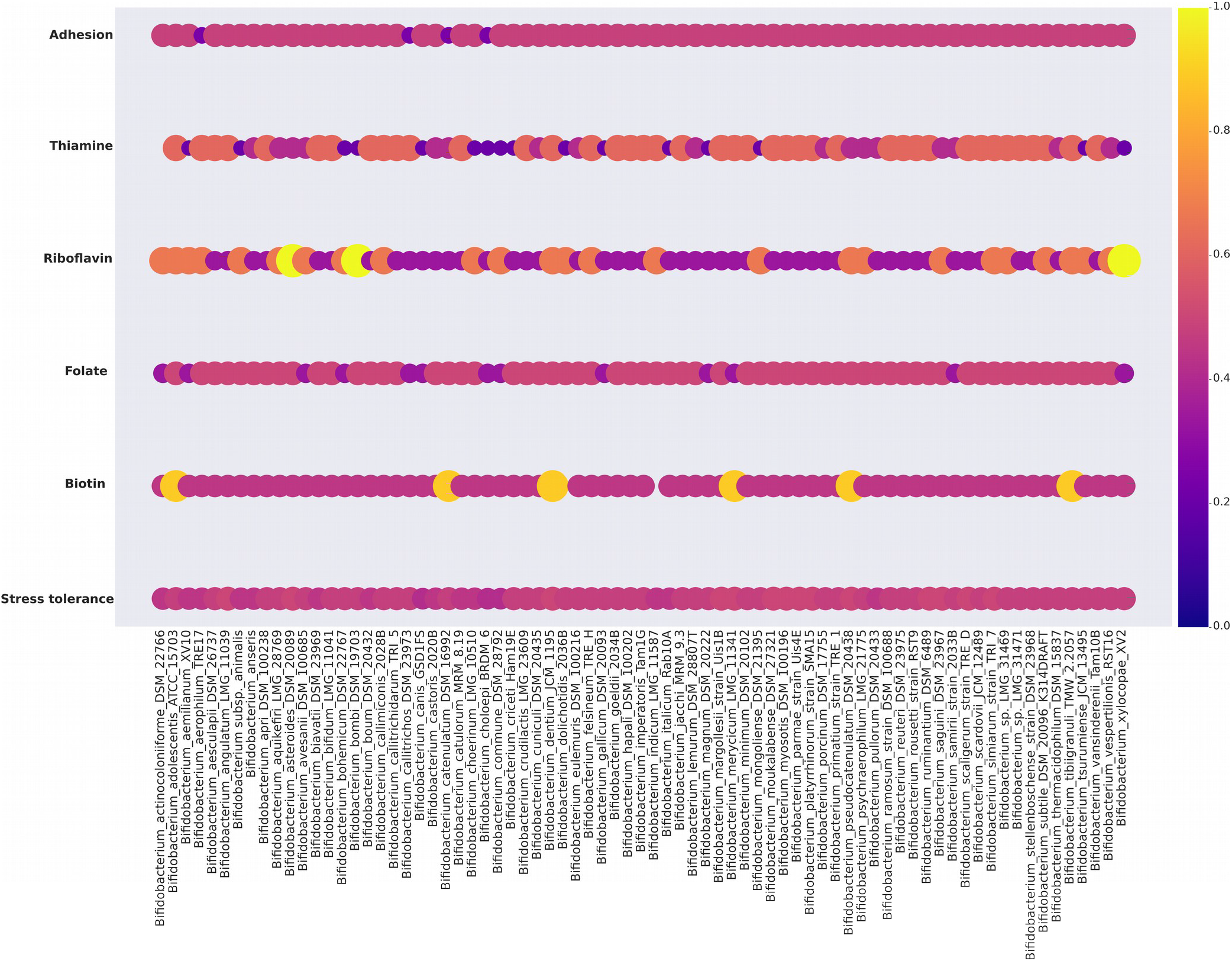
Relative abundance of genes involve in probiotic functions among the type species of bifidobacerium genus.

